# A benchmarking study on virtual ligand screening against homology models of human GPCRs

**DOI:** 10.1101/284075

**Authors:** Victor Jun Yu Lim, Weina Du, Yu Zong Chen, Hao Fan

## Abstract

G-protein-coupled receptor (GPCR) is an important target class of proteins for drug discovery, with over 27% of FDA-approved drugs targeting GPCRs. However, being a membrane protein, it is difficult to obtain the 3D crystal structures of GPCRs for virtual screening of ligands by molecular docking. Thus, we evaluated the virtual screening performance of homology models of human GPCRs with respect to the corresponding crystal structures. Among the 19 GPCRs involved in this study, we observed that 10 GPCRs have homology models that have better or comparable performance with respect to the corresponding X-ray structures, making homology models a viable choice for virtual screening. For a small subset of GPCRs, we also explored how certain methods like consensus enrichment and sidechain perturbation affect the utility of homology models in virtual screening, as well as the selectivity between agonists and antagonists. Most notably, consensus enrichment across multiple homology models often yields results comparable to the best performing model, suggesting that ligand candidates predicted with consensus scores from multiple models can be the optimal option in practical applications where the performance of each model cannot be estimated.

## Introduction

GPCRs, also commonly known as seven-transmembrane domain receptors, are responsible for many of our physiological responses and activities, including responses to hormone, neurotransmitter and even environmental stimulants such as taste, smell and vision^1^. GPCRs represent the largest and most successful class of druggable targets in the human genome, with over 27% of FDA-approved drugs targeting approximately 60 out of the total over 800 GPCRs^2–3^. However, majority of human GPCRs have not yet been explored in drug discovery. Thus, tremendous efforts are now being made to exploit the remaining receptors.

Virtual screening against GPCR structures, is a computational approach to identify ligands and is often a rewarding effort^4–7^. However, the utility of this approach depends on the availability of GPCR structures determined by X-ray crystallography. Despite the recent bloom in the number of GPCR structures^8–11^, the number of human GPCRs with X-ray structures is only 25 out of a total of 842^12^, 3% of the entire human GPCR proteome. This limitation highly restricts the potential of protein structure-based approach such as virtual screening in drug discovery of GPCRs.

One way to overcome this limitation is through 3D structure prediction of GPCRs using the homology modelling technique, based on known structures of closely related proteins (templates). As seen from other classes of proteins, homology models used for virtual screening can be successful^13^. With homology modelling, it is possible to model up to 30% of all human GPCRs at the minimum of 20% sequence identity, and up to 10% of human GPCRs at the minimum of 30% sequence identity. This is a great increase compared to the current 3% of known human GPCR structures.

It has been shown that it is possible for homology models of GPCRs to identify potential ligands^14^. There have been a number of studies aiming to evaluate the performance of homology models of GPCRs for *in-silico* drug screening^14–21^. For example, Tang et al. studied beta-2 adrenoreceptor (ADRB2) and showed that some homology models could exceed crystal structures in ligand virtual screening^22^. However, all of them only focus on one or a few GPCRs and their homology models, which do not provide a complete picture about the capability of homology model based methods for virtual screening of GPCR ligands. Thus, we aim to do a large-scale evaluation on the performance of homology models of human GPCRs with respect to that of the corresponding X-ray structures. For each GPCR target, we selected templates from three ranges of sequence identity, to investigate how target-template sequence identity affects the performance of homology models of GPCRs in terms of ligand prediction.

Furthermore, to gain further insights on virtual ligand screening of GPCRs, we are interested in the following questions. First, since there are usually multiple homology models present for a receptor, we wanted to know if the consensus of ligand enrichment across multiple models can have an impact on the performance of virtual screening. Second, since protein sidechains are flexible, we intended to explore that flexibility with respect to different ligands and see how they affect the virtual screening. Third, we investigated if virtual screening methods have any inclination to select for higher affinity ligands over lower affinity ligands, given the fact that higher affinity ligands are more likely potential drug leads or chemical probes than lower affinity ligands. Lastly, we were interested to know if the modelled structures have selectivity for agonists or antagonists. Agonists and antagonists have different biological and pharmacological implications on the cell signalling, disease processes and therapeutic actions. Therefore, it is essential to differentiate them during virtual screening.

## Materials & Methods

### GPCR targets and template selection

We started from the ligand-bound (holo) crystal structures and ligands of 24 GPCRs that were used in a benchmarking study of GPCR crystal structures^23^. Among these 24 GPCRs, ACM3, ADRB1, OPRM and OPRD were not included in this study because X-ray structures were not available for human proteins, neither were the majority of ligands in the database. 20 human GPCR sequences were obtained from UniProt^24^ database. To find templates for each GPCR, their protein sequences were BLAST^25^ aligned to the sequences of the entries of the PDB database. Up to 3 templates were selected for each protein for homology modelling in the following range of sequence identity to the target: 20-30%, 30-50% and 50-80%. The range of sequence identity were selected to correspond to low, medium where the accuracy is equivalent to a low resolution X-ray structure and high where there is little need for manual adjustment to alignment for a reliable model^26^. The template would only be considered if it has an X-ray resolution of at least 3.4 Å. Furthermore, within each sequence identity range, we selected the template that has the closest distance to the target in the GPCR phylogenetic tree^27^, so that the templates more likely have similar structures as the targets and result in homology models of the latter of as high quality as possible. Out of the 20 GPCRs, SMO do not have template available under our criteria and were dropped. The list of target-template pairs is shown in Table 1, a total of 38 target-template pairs for 19 GPCRs.

**Table 1.** GPCR targets and their templates for homology modelling

| Table 1. GPCR targets and their templates for homology modelling |  |  |  |  |  |  |  |
| --- | --- | --- | --- | --- | --- | --- | --- |
| Receptor | PDBID | Resolution (Å) | Sequence identity range % | Sequence identity (%) | Template PDBID | Template protein | Resolution (Å) |
| 5HT1B | 4IAR | 2.7 | 20-30 | 27 | 4IB4 | 5HT2B | 2.7 |
|  |  |  | 30-50 | 36 | 3PBL | DRD3 | 2.9 |
| 5HT2B | 4IB4 | 2.7 | 20-30 | 27 | 4IAR | 5HT1B | 2.7 |
|  |  |  | 30-50 | 36 | 5DSG | ACM4 | 2.6 |
| AA2AR | 4EIY | 1.8 | 20-30 | 25 | 3UON | ACM2 | 3.0 |
|  |  |  | 30-50 | 36 | 2VT4 | ADRB1 | 2.7 |
| ACM2 | 3UON | 3.0 | 20-30 | 25 | 3PBL | DRD3 | 2.9 |
|  |  |  | 30-50 | 44 | 5CXV | ACM1 | 2.7 |
|  |  |  | 50-80 | 56 | 5DSG | ACM4 | 2.6 |
| ADRB2 | 2RH1 | 2.4 | 20-30 | 27 | 4IB4 | 5HT2V | 2.7 |
|  |  |  | 30-50 | 36 | 3PBL | DRD3 | 2.9 |
|  |  |  | 50-80 | 58 | 2Y00 | ADRB1 | 2.5 |
| CCR5 | 4MBS | 2.7 | 20-30 | 27 | 5C1M | OPRM | 2.1 |
|  |  |  | 30-50 | 34 | 3ODU | CXCR4 | 2.5 |
| CRFR1 | 4K5Y | 3.0 | 30-50 | 37 | 5EE7 | GCCR | 2.5 |
| CXCR4 | 3ODU | 2.5 | 20-30 | 28 | 4MBS | CCR5 | 2.7 |
|  |  |  | 30-50 | 36 | 4ZUD | AT1R | 2.8 |
| DRD3 | 3PBL | 2.9 | 20-30 | 25 | 4EIY | AA2AR | 1.8 |
|  |  |  | 30-50 | 41 | 3V2W | S1PR1 | 3.4 |
| GPR40 | 4PHU | 2.3 | 20-30 | 27 | 3VW7 | PAR1 | 2.2 |
| HRH1 | 3RZE | 3.1 | 20-30 | 27 | 2RH1 | ADRB2 | 2.4 |
|  |  |  | 30-50 | 33 | 4IAR | 5HT1B | 2.7 |
| MGLUR1 | 4OR2 | 2.8 | 50-80 | 75 | 4OO9 | MGLUR5 | 2.6 |
| MGLUR5 | 4OO9 | 2.6 | 50-80 | 79 | 4OR2 | MGLUR1 | 2.8 |
| OPRK | 4DJH | 2.9 | 20-30 | 27 | 4MBS | CCR5 | 2.7 |
|  |  |  | 30-50 | 34 | 4ZUD | AT1R | 2.8 |
|  |  |  | 50-80 | 69 | 5C1M | OPRM | 2.1 |
| OPRX | 4EA3 | 3.0 | 20-30 | 24 | 4MBS | CCR5 | 2.7 |
|  |  |  | 30-50 | 32 | 3ODU | CXCR4 | 2.5 |
|  |  |  | 50-80 | 61 | 5C1M | OPRM | 2.1 |
| OX2R | 4S0V | 2.5 | 20-30 | 25 | 4XEE | NTSR1 | 2.9 |
|  |  |  | 30-50 | 30 | 2YDO | AA2AR | 3.0 |
|  |  |  | 50-80 | 75 | 4ZJ8 | OX1R | 2.8 |
| P2Y12 | 4NTJ | 2.6 | 20-30 | 25 | 2VT4 | ADRB1 | 2.7 |
|  |  |  | 30-50 | 30 | 4EJ4 | OPRD | 3.4 |
| PAR1 | 3VW7 | 2.2 | 20-30 | 25 | 4N6H | OPRD | 1.8 |
| S1PR1 | 3V2Y | 2.8 | 20-30 | 25 | 3OAX | RHO | 2.6 |
|  |  |  | 30-50 | 36 | 4Z34 | LPAR1 | 3.0 |

### Homology modelling

For each of the target-template pairs in Table 1, the sequence alignment for homology modelling was obtained using the profile-profile method^28^. The profiles of the target and template were obtained by performing BLAST sequence alignment to the non-redundant database and the top 500 sequences were used as the profile. The profiles of the target and template were aligned using MUSCLE v3.8^29^. Homology models of each target were generated using MODELLERv9.13^30^, 500 models were generated for each target-template pair. DOPE^31^ scoring function was then used to score the models and the best scoring model was used for virtual ligand screening.

### Virtual ligand screening

The ligands of the 19 GPCRs were obtained from Weiss et al.^23^, as mentioned above. Property-matching decoys were generated for each GPCR using the Database of Useful Decoys, Enhanced (DUD-E)^32^ approach based on the compounds from the ZINC^33^ database. The ligands and decoys were then docked to the X-ray structure of each target and corresponding homology models using DOCK v3.6^34^. The sampling space during docking was determined by the ligand binding site in the holo X-ray structures, and homology models were superposed on the corresponding X-ray structures. The docked compounds were ranked by the docking energy function that is the sum of van der Waals, Poisson-Boltzmann electrostatic, and ligand desolvation penalty terms.

### Evaluation of virtual screening results

The accuracy of the virtual screening was evaluated using enrichment factor (EF) and logAUC as described by Fan et al.^35^ and Mysinger et al.^32^. The EF measures the ratio of known ligands that was found among the top scoring compounds as compared to random selection and is defined as follows:

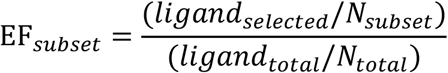

where *ligand_total_* is the total number of ligand in the database of *N_total_* compound and *ligand_selected_* is the number of ligands found in the subset of *N_subset_* top scoring compounds.

However, EF only accounts for a fixed subset of all docked compound (eg. 1% or 5%). Therefore, another measure that incorporates information considering all the different subsets was introduced. An enrichment curve is a plot of the percentage of actual ligand found (y-axis) within the top ranked subsets of the database against the entire ranked database (x-axis on logarithm scale). LogAUC (logarithmic Area Under Curve) represents the area under the enrichment curve with more emphasis given to early enrichment by using logarithmic scale for the x-axis. LogAUC is defined as follow:

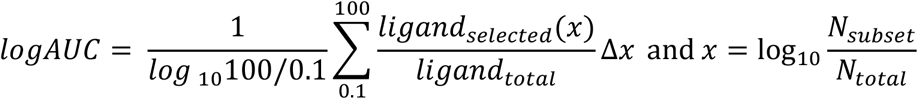

where Δx is 0.1. A random selection of ligands from the database will yield a logAUC value of 14.5 while a mediocre selection picking twice as many ligands than random yields a logAUC of 24.5. When comparing two structures/models, it is considered that their individual enrichment to be of significant difference from each other if there is a difference of at least 3 logAUC unit, otherwise, the enrichment values are comparable.

### Consensus over multiple homology models

For a subset of receptors from the 19 GPCRs, namely AA2AR and ADRB2 that have multiple crystal structures determined in complex with both agonists and antagonists, further investigation was done by looking at multiple X-ray structures and homology models that each of these models is based on a different template. These structures/models were selected to have different ligands and a balance ratio of agonist-bound to antagonist-bound X-ray structures and templates. This is to ensure that our target and template structures cover a larger range of conformations available for the active sites for protein-ligand complementarity. The additional target and template X-ray structures are shown in the supporting information (Table S1).

For each target structure or homology model, its individual enrichment score was computed, as well as a consensus enrichment score for each target protein. The consensus score was calculated by taking the best docking energy for each compound across all structures/models and then used in ranking^35^.

### Alternative sidechain conformations

For the same subset of 2 GPCRs (AA2AR and ADRB2), sidechains in the X-ray structures and homology models listed in Table 1 were rebuilt by sidechain prediction methods. The two prediction programs Protein Local Optimization Program (PLOP)^36^ and SCWRL4^37^ were used. For X-ray structures of each target, sidechains were predicted in the presence of the cognate ligand in the active site. For each homology model, multiple ligands of that receptor were docked to that homology model and then sidechains were predicted with each docked ligand, yielding multiple models. After sidechain reconstruction, the resulted structures/models were subjected to another round of virtual screening, followed by ligand enrichment calculations. For homology models, since there are multiple predicted models from different docked ligands, we calculated the consensus enrichment using the same method described in “Consensus over multiple homology models”.

### High affinity ligands

Ligands for AA2AR and ADRB2 were filtered by a Ki or Kd cut-off value of 10nM. The cut-off was selected so that a good number of high affinity ligands can be found in our ligand set. Then the enrichment score was calculated by considering only ligands with better Ki or Kd values as true ligands while the rest in the database as decoys. For AA2AR, 70 out of 208 original ligands have Ki/Kd values lesser than 10nM, and for ADRB2, 27 out of 207. The ChEMBL IDs of these high affinity ligands are in the supporting information (Table S2). The high affinity ligands and the rest compounds were then docked to the X-ray structures and homology models described in “Consensus over multiple homology models”.

### Agonist/Antagonist selectivity

For AA2AR and ADRB2, their agonists and antagonists were manually curated from ChEMBL^38^ database. For AA2AR, there are 9 annotated agonists and 9 antagonists and for ADRB2, 11 agonists and 11 antagonists. The agonists and antagonists were then docked to the X-ray structures and homology models described in “Consensus over multiple homology models”. T-scores were calculated based on the docking energies of the agonists and antagonists to see if there was significant difference between them. The ChEMBL IDs of the agonists and antagonists for AA2AR and ADRB2 are in the supporting information (Table S3).

## Results and Discussions

### Cognate docking and geometry assessment

To examine if the docking method is robust enough to be used for this benchmarking study, we docked the cognate ligand back into the X-ray structure of each target, and calculated the root mean-square deviation (RMSD) between the ligand docking pose and the ligand crystal structure. This examines whether docking method used can reproduce the crystal conformation accurately (RMSD value < 3 Å, near-native) The RMSD values are shown in Table 2.

**Table 2.** Structural deviations of docking poses from crystal structures of cognate ligands

|  |  |  |  |
| --- | --- | --- | --- |
| Receptor | PDBID | RMSD (best energy) (Å) | Lowest RMSD (Å) |
| 5HT1B | 4IAR | **0.56** | **0.56** |
| 5HT2B | 4IB4 | **0.53** | **0.53** |
| AA2AR | 4EIY | 3.73 | **1.67** |
| ACM2 | 3UON | **0.81** | **0.78** |
| ADRB2 | 2RH1 | **0.75** | **0.75** |
| CCR5 | 4MBS | **1.07** | **1.07** |
| CRFR1 | 4K5Y | **0.50** | **0.50** |
| CXCR4 | 3ODU | **1.13** | **1.13** |
| DRD3 | 3PBL | 9.66 | **1.64** |
| GPR40 | 4PHU | **2.06** | **1.39** |
| HRH1 | 3RZE | **1.25** | **0.98** |
| MGLUR1 | 4OR2 | **1.23** | **1.23** |
| MGLUR5 | 4OO9 | **2.40** | **2.30** |
| OPRK | 4DJH | 5.55 | 5.55 |
| OPRX | 4EA3 | **2.89** | **0.87** |
| OX2R | 4S0V | 6.35 | **2.11** |
| P2Y12 | 4NTJ | 11.18 | 4.57 |
| PAR1 | 3VW7 | **0.40** | **0.40** |
| S1PR1 | 3V2Y | **2.22** | **1.59** |
| **RMSD (best energy)**, the RMSD value of the ligand docking pose with the best docking energy. **Lowest RMSD**, the best RMSD value among all the docking poses sampled. RMSD value in Bold indicates that we reproduce the crystal conformation accurately (RMSD <3 Å) by docking. |  |  |  |

In 14 out of the 19 GPCR X-ray structures, the ligand crystal structures were accurately reproduced by the best scoring docking pose, as seen in Figure 1A and 1B. For the rest of the 5 GPCRs, higher RMSD values can be attributed to two possible reasons: imperfect scoring and insufficient sampling. In the cases of AA2AR, DRD3, and OX2R, the best RMSD value among all the docking poses sampled is below 3 Å while the docking pose with the best energy is not. For example, the docking energy function was not able to rank the near-native conformation the best for DRD3 (Figure 1C), which is likely attributed to the under-estimation of hydrophobic interactions between the crystal ligand and the inner-faces of the binding pocket. While for OPRK and P2Y12, the lowest RMSD is above 3 Å. This indicates that the sampling is insufficient to approach the ligand crystal structure. For P2Y12 (Figure 1D), the insufficient sampling could be partially caused by a higher number (8) of rotatable bonds in the crystal ligand in P2Y12 with respect to that in the crystal ligand in PAR1 (Figure 1A) and in ADRB2 (Figure 1B) containing 6 rotatable bonds. Another possible reason could be that two crystal water molecules that form hydrogen bonds with the sulfone group of the crystal ligand, were removed during the docking process of P2Y12.

**Figure 1.**
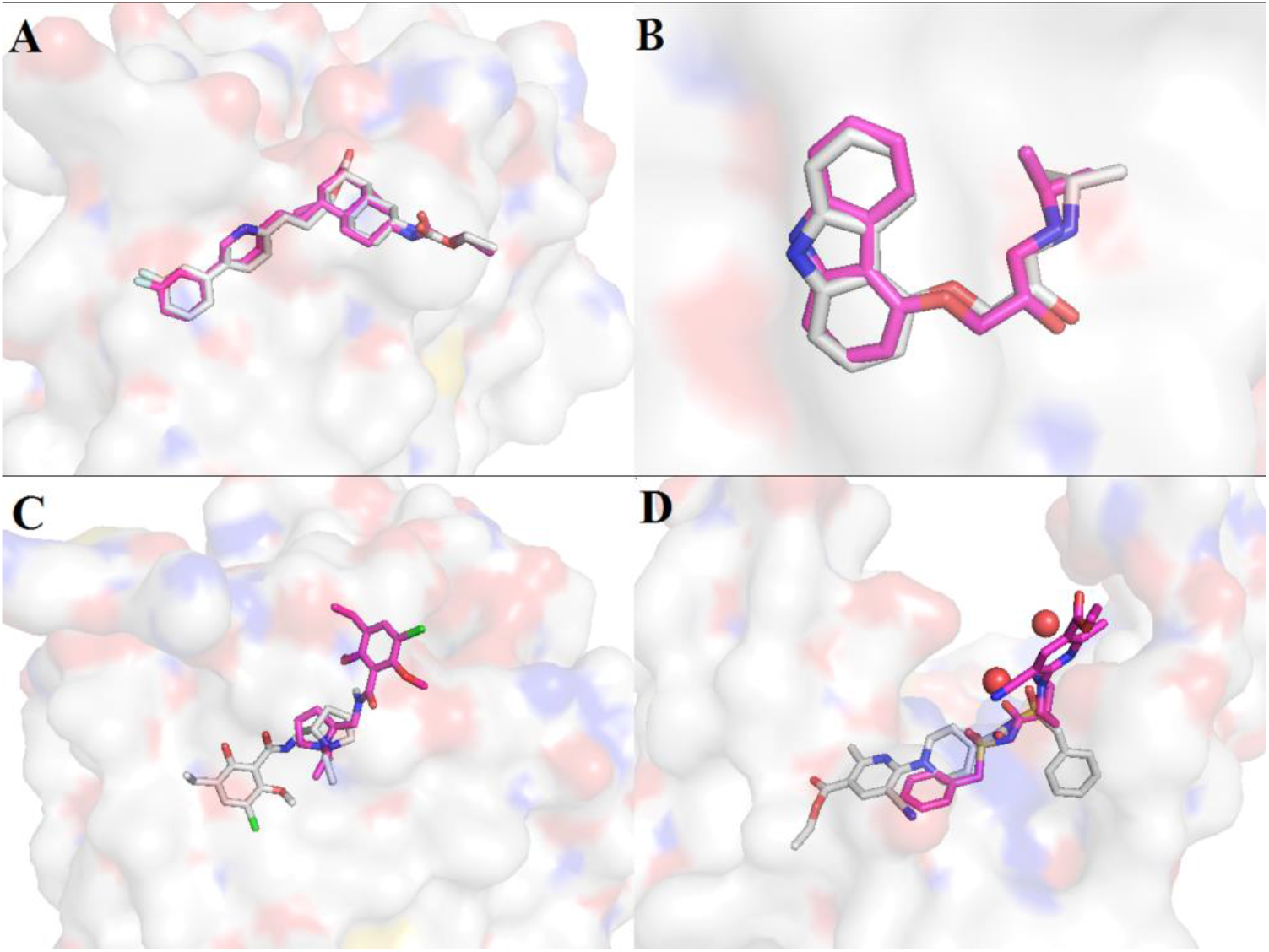
Docking poses (colored in magenta) of cognate ligands compared to the corresponding crystal structures (colored in white). (A) Top left, PAR1, PDBID 3VW7. (B) Top right, ADRB2, PDBID 2RH1. (C) Bottom left, DRD3, PDBID 3PBL. (D) Bottom right, P2Y12, PDBID 4NTJ.

### Virtual screening and ligand enrichment

After docking all the ligands and decoys to the X-ray structures and homology models, we plotted their enrichment curves and calculated their logAUC and EF_1_ values. The results are shown in Figure 2 and Table 3. Out of the 38 homology models of the 19 GPCRs, 23 (60.5%), 9 (23.7%), and 6 (15.8%) of them showed worse, comparable, and better virtual screening performance, respectively, with respect to that from corresponding X-ray structures. The results are expected as homology models are predicted and should contain structural errors. Nonetheless, our study clearly showed that it is still possible for homology models to be useful in ligand recognition. This is indicated by the fact that among the 19 GPCRs, 10 of them (5HT2B, ADRB2, CXCR4, HRH1, MGLUR1, MGLUR5, OPRK, OX2R, P2Y12, S1PR1) have at least one homology model performing better than or comparable to their X-ray structures.

**Figure 2.**
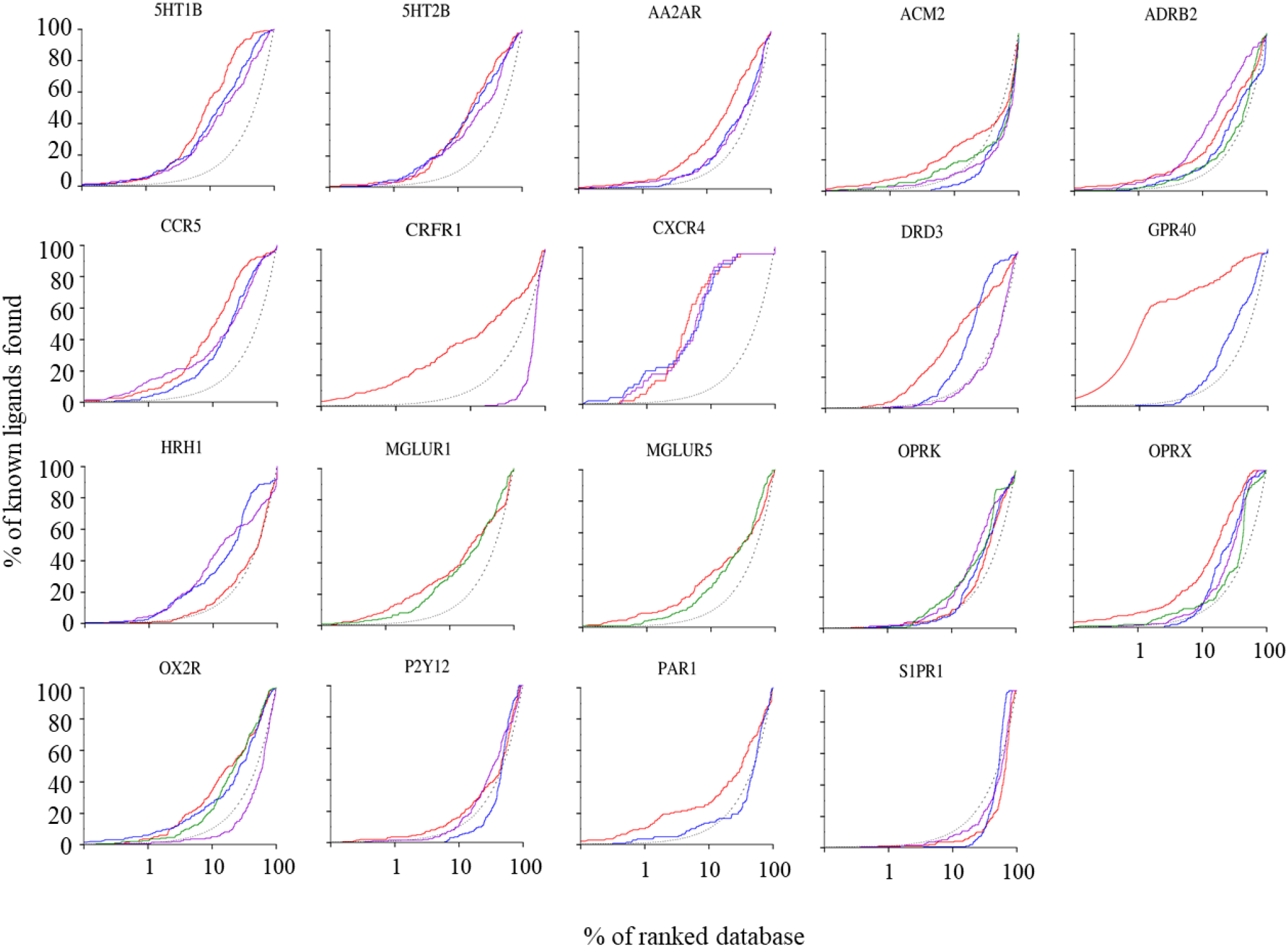
Enrichment plots for the 19 GPCRs, including the X-ray structures (red), homology models of 20-30% sequence identity (blue), homology models of 30-50% sequence identity (purple), homology models of 50-80% sequence identity (green), and random selection (dotted line).

**Table 3.**
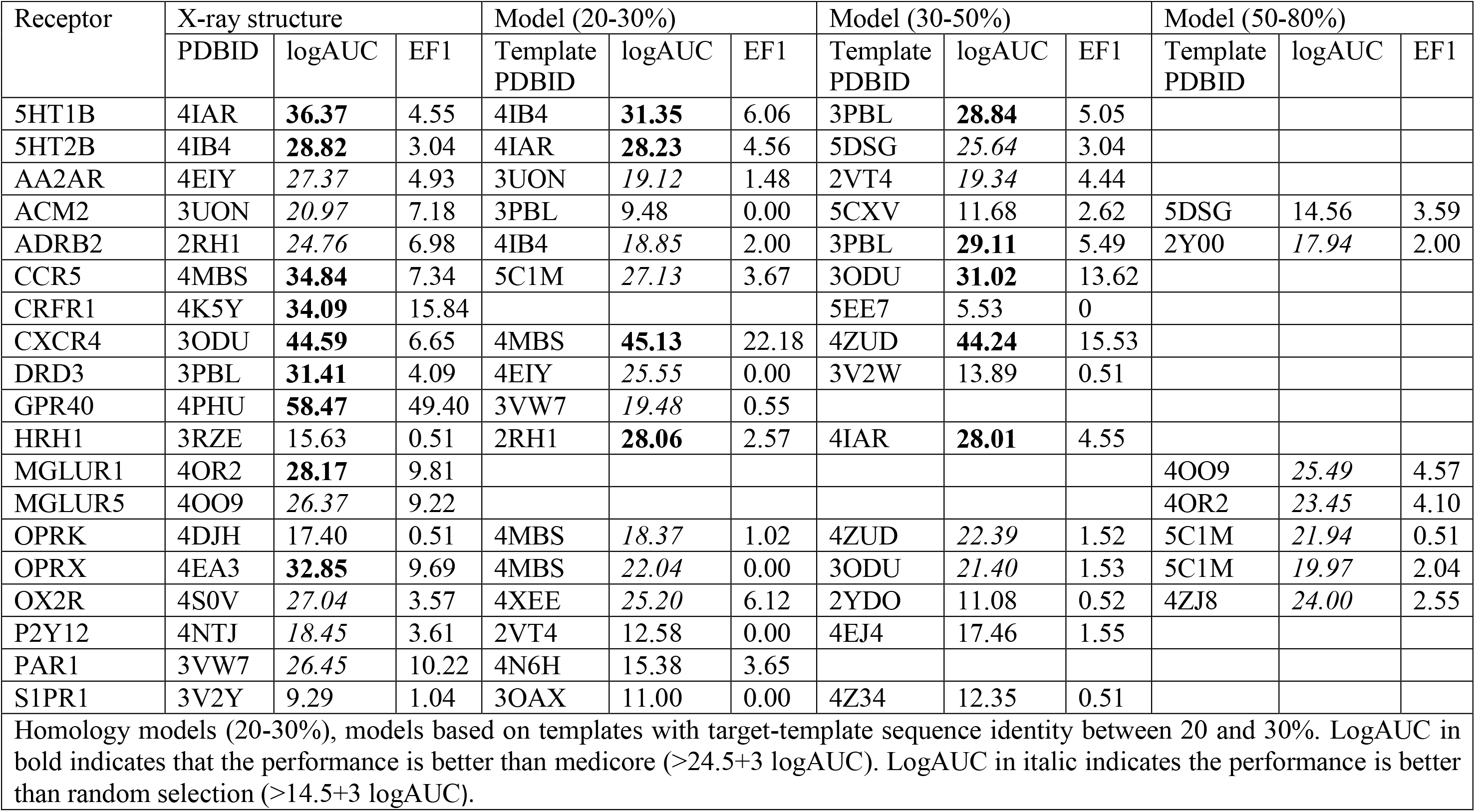
Ligand enrichment of X-ray structures and homology models

| Table 3. Ligand enrichment of X-ray structures and homology models |  |  |  |  |  |  |  |  |  |  |  |  |
| --- | --- | --- | --- | --- | --- | --- | --- | --- | --- | --- | --- | --- |
| Receptor | X-ray structure |  |  | Model (20-30%) |  |  | Model (30-50%) |  |  | Model (50-80%) |  |  |
|  | PDBID | logAUC | EF1 | Template PDBID | logAUC | EF1 | Template PDBID | logAUC | EF1 | Template PDBID | logAUC | EF1 |
| 5HT1B | 4IAR | **36.37** | 4.55 | 4IB4 | **31.35** | 6.06 | 3PBL | **28.84** | 5.05 |  |  |  |
| 5HT2B | 4IB4 | **28.82** | 3.04 | 4IAR | **28.23** | 4.56 | 5DSG | *25.64* | 3.04 |  |  |  |
| AA2AR | 4EIY | *27.37* | 4.93 | 3UON | *19.12* | 1.48 | 2VT4 | *19.34* | 4.44 |  |  |  |
| ACM2 | 3UON | *20.97* | 7.18 | 3PBL | *9.48* | 0.00 | 5CXV | *11.68* | 2.62 | 5DSG | *14.56* | 3.59 |
| ADRB2 | 2RH1 | *24.76* | 6.98 | 4IB4 | *18.85* | 2.00 | 3PBL | **29.11** | 5.49 | 2Y00 | *17.94* | 2.00 |
| CCR5 | 4MBS | **34.84** | 7.34 | 5C1M | *27.13* | 3.67 | 3ODU | **31.02** | 13.62 |  |  |  |
| CRFR1 | 4K5Y | **34.09** | 15.84 |  |  |  | 5EE7 | *5.53* | 0 |  |  |  |
| CXCR4 | 3ODU | **44.59** | 6.65 | 4MBS | **45.13** | 22.18 | 4ZUD | **44.24** | 15.53 |  |  |  |
| DRD3 | 3PBL | **31.41** | 4.09 | 4EIY | *25.55* | 0.00 | 3V2W | *13.89* | 0.51 |  |  |  |
| GPR40 | 4PHU | **58.47** | 49.40 | 3VW7 | *19.48* | 0.55 |  |  |  |  |  |  |
| HRH1 | 3RZE | *15.63* | 0.51 | 2RH1 | **28.06** | 2.57 | 4IAR | **28.01** | 4.55 |  |  |  |
| MGLUR1 | 4OR2 | **28.17** | 9.81 |  |  |  |  |  |  | 4OO9 | *25.49* | 4.57 |
| MGLUR5 | 4OO9 | *26.37* | 9.22 |  |  |  |  |  |  | 4OR2 | *23.45* | 4.10 |
| OPRK | 4DJH | *17.40* | 0.51 | 4MBS | *18.37* | 1.02 | 4ZUD | *22.39* | 1.52 | 5C1M | *21.94* | 0.51 |
| OPRX | 4EA3 | **32.85** | 9.69 | 4MBS | *22.04* | 0.00 | 3ODU | *21.40* | 1.53 | 5C1M | *19.97* | 2.04 |
| OX2R | 4S0V | *27.04* | 3.57 | 4XEE | *25.20* | 6.12 | 2YDO | *11.08* | 0.52 | 4ZJ8 | *24.00* | 2.55 |
| P2Y12 | 4NTJ | *18.45* | 3.61 | 2VT4 | *12.58* | 0.00 | 4EJ4 | *17.46* | 1.55 |  |  |  |
| PAR1 | 3VW7 | *26.45* | 10.22 | 4N6H | *15.38* | 3.65 |  |  |  |  |  |  |
| S1PR1 | 3V2Y | *9.29* | 1.04 | 3OAX | *11.00* | 0.00 | 4Z34 | *12.35* | 0.51 |  |  |  |
| Homology models (20-30%), models based on templates with target-template sequence identity between 20 and 30%. LogAUC in bold indicates that the performance is better than medicore (>24.5+3 logAUC). LogAUC in italic indicates the performance is better than random selection (>14.5+3 logAUC). |  |  |  |  |  |  |  |  |  |  |  |  |

Although it is not surprising that over 50% of homology models tested do not perform as well as the X-ray structures, there is still a need to evaluate the general utility of the models for virtual screening, as many GPCRs have no X-ray structure. An analysis of the percentage of models that perform 3 logAUC unit better than random selection (logAUC > (14.5+3)) and value-add to ligand screening was performed (Table 4). As seen from Table 4, homology models built with sequence identity 50-80% to targets have the highest chance (84%) to value-add to virtual screening, which is comparable to that of X-ray structures (83%). This is likely a consequence that higher sequence identity templates are more similar in structures to the targets and thus the models generated are likely closer to the true structures. Noticeably, models with sequence identity 20-30% and 30-50% to targets are still useful, having significant probability (75% and 60%) to value-add to virtual screening, therefore homology models can be used when there are no alternatives. It should also be noted that X-ray structures have the highest probability (47%) of having a good enrichment-selecting more than twice the number of ligands than random (logAUC > 24.5+3).

**Table 4.** Value added to virtual ligand screening by homology models

| **Table 4. Value added to virtual ligand screening by homology models** |  |  |  |  |  |  |
| --- | --- | --- | --- | --- | --- | --- |
| Structure type | Sequence Identity | Mediocre ligand enrichment | Good ligand enrichment | Value added models/ structures | Total number of models/ structures | Percent (%) of value added models/ structures |
|  |  | 17.5-27.5 logAUC | >27.5 logAUC | >17.5 logAUC |  |  |
| Homology model | 20-30% | 8 | 4 | 12 | 16 | 75 |
|  | 30-50% | 5 | 4 | 9 | 15 | 60 |
|  | 50-80% | 6 | 0 | 6 | 7 | 86 |
|  | Total | 19 | 8 | 27 | 37 | 73 |
| X-ray structure |  | 7 | 9 | 16 | 19 | 84 |
| **Mediocre ligand enrichment**, perform better than random selection but does not select more than twice as many ligands in any given subset ($14.5+3 < \text{logAUC} < 24.5+3$). **Good ligand enrichment**, select for more than twice as ligands in any given subset ($\text{logAUC} > 24.5+3$) | | | | | | |

We note that there is an over representation of class A GPCRs. This is because at the time this paper is written, class A GPCRs were more intensively studied, with more structures available. Out of the 19 GPCRs benchmarked, 16 of them belongs to class A, 1 class B (CRFR1) and 2 class C (MGLUR1 and MGLUR2). However, it is observed that the GPCRs in class B and C shows similar performance to class A GPCRs, thus it is unlikely that this poses a problem in the generalisability of the results to other classes of GPCRs.

Also, for MGLUR1 and MGLUR5, their X-ray structures were solved with their negative allosteric modulator instead of orthosteric agonist or antagonist. However, the allosteric site overlaps with the orthosteric site of class A receptors^39^ and it is noted that antagonist for MGLUR1 are mainly designed for its allosteric site within the 7 transmembrane domain as the targeting the orthosteric site shows poor selectivity and antagonist^40^. This is demonstrated in our results that both X-ray structures and homology models of the two MGLUR receptors show reasonable enrichment of database ligands (logAUC > 17.5). Therefore, docking to the allosteric site of both MGLUR receptors is justifiable for our study.

### Consensus over multiple homology models

For the subset of 2 GPCRs, AA2AR and ADRB2, their individual and consensus logAUC were calculated (Table 5). The results shown are consistent with the conclusion above that homology models based on higher sequence identity templates can have a higher chance to value-add (logAUC > 14.5+3) to virtual screening. For this subset, homology models based on 20-30% and 30-50% sequence identity templates have 62% and 75% chance of value-add to ligand screening, while both X-ray structures and homology models based on 50-80% sequence identity templates have 100% chance to value-add.

**Table 5.**
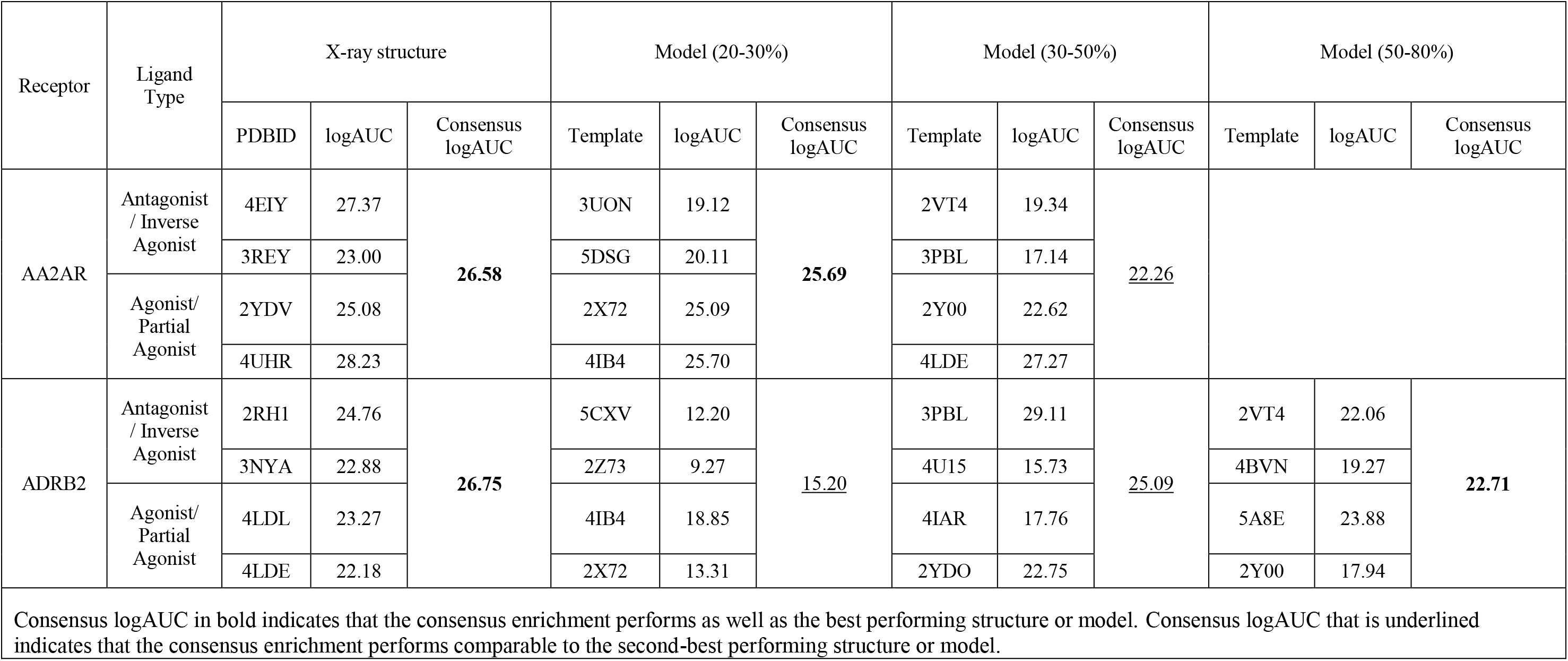
Ligand enrichment of multiple X-ray structures and homology models for AA2AR and ADRB2

More importantly, for the consensus results over multiple structures/models, the majority (57%) of all the consensus logAUC values are comparable to the best performing structures/models, and the rest are comparable to the second best. This is consistent with the observation by Kelemen et al. that consensus scoring over multiple crystal structures works well for ligands selections in class A aminergic GPCRs^41^. This is very promising, as simply by taking the consensus score for each compound over multiple homology models, it is possible to approach the ligand recognition ability of the best performing model that would often be difficult to select when crystal structure has not been solved. This could be because there is always certain conformation of the binding site complementing the ligands but not the decoys regardless of the sidechains conformation. This may lead to an advantage for the ligands to be better recognized using consensus scoring than decoys.

### Alternative sidechain conformations

Sidechains in each X-ray structure were predicted together with the cognate ligand and the modified structure was evaluated in virtual screening. For X-ray structures of AA2AR and ADRB2, there is not significant changes in the ligand enrichment (logAUC), as shown in the supporting information (Table S4). Although preliminary, it seems there is no drawback to regenerate sidechain conformations in GPCR X-ray structures for virtual screening. A likely explanation is that X-ray crystallography only capture one snapshot of the conformational ensemble of a protein. Thus, an alternative conformation generated by rebuilding the sidechains could also offer similar complementarity for ligands in the database.

The performance of the homology models after sidechain reconstruction with docked ligands is evaluated (Table 6). First, we evaluate the results by checking for significant difference (3 logAUC unit difference) between the consensus logAUC of the modified models and that of the original model. Using SCWRL4, 4 out of 5 sets of modified models showed significantly improved consensus logAUC values while using PLOP, only 1 sets of the models showed significant improvement. The difference in performance is likely due to different parameters used to run the programs. For SCWRL4, all sidechains were regenerated while for PLOP, only the binding site residues were predicted. This might restrict certain conformation of the protein due to steric hindrance from some residues and thus limit the performance of PLOP. Regarding the best logAUC, it is improved in all cases using SCWRL4 while 4 out of 5 cases (80%) using PLOP. This is promising because if there is knowledge on known ligands of the target protein, it is possible to perform sidechain sampling to generate multiple models that will be selected for virtual screening.

**Table 6.** Ligand enrichment of homology models with predicted sidechains

|  |  |  |  |  |  |  |
| --- | --- | --- | --- | --- | --- | --- |
| Receptor | Sequence Identity (%) | Template PDBID | Model logAUC | Prediction program | Consensus logAUC | Best logAUC |
| AA2AR | 25 | 3UON | 19.12 | SCWRL | **23.83** | **29.88** |
|  |  |  |  | PLOP | 19.91 | **26.98** |
|  | 36 | 2VT4 | 19.34 | SCWRL | 20.30 | **25.32** |
|  |  |  |  | PLOP | 19.21 | **23.10** |
| ADRB2 | 27 | 4IB4 | 18.85 | SCWRL | **23.34** | **29.84** |
|  |  |  |  | PLOP | **23.87** | **26.39** |
|  | 36 | 3PBL | 29.11 | SCWRL | **34.16** | **40.30** |
|  |  |  |  | PLOP | 18.48 | 23.29 |
|  | 58 | 2Y00 | 17.94 | SCWRL | **28.97** | **29.01** |
|  |  |  |  | PLOP | 19.35 | **26.18** |
| **Best logAUC**, refers to the highest logAUC attained by the models with sidechains reconstructed. Consensus logAUC in bold indicates that the consensus logAUC from multiple modified models is significantly better than that of the original model ($\Delta\text{logAUC}>3$). | | | | | | |

### High affinity ligands

For high affinity ligands, we compared the logAUC of their enrichments with those of all ligands (Table 7). For AA2AR, no structures and models show significantly selectivity for high affinity ligands compared to all the ligands. The ligand docking energy distribution from one AA2AR structure (PDBID 4EIY) is shown as an example (Figure S1) where high affinity ligands and low affinity ligands have similar distribution in terms of docking energies. For ADRB2, there are structures (2 out of 4) and models (5 out of 11) that seem to be more selective for high affinity ligands. This is likely due to the small number (27) of high affinity ligands for ADRB2, thus the observation may not be statistically significant. For the X-ray structures of the 2 GPCRs, the chemical similarity measured by the Tanimoto coefficient (Tc)^42^ calculated using ECPF4 between the cognate ligands and database ligands were calculated, showing no significant similarity (Tc>0.4) between the cognate ligands and the database ligands (neither the high affinity ligands nor the low affinity ligands). This likely explains why there is no better selectivity over high affinity ligands. Another possible reason for the lack of selectivity for high affinity ligands is that the scoring function is not accurate enough to properly measure the ligand binding affinity. As seen from cognate ligand docking, there are certain interactions such as hydrophobic interaction between ligand and protein that is underestimated with the current docking energy.

**Table 7.** Ligand enrichment of X-ray structures and homology models for high affinity ligands

|  |  |  |  |  |  |  |  |
| --- | --- | --- | --- | --- | --- | --- | --- |
| Receptor | X-ray structure |  |  | Homology model |  |  |  |
|  | PDBID | LogAUC for all the ligands | LogAUC for high affinity ligands | Target-template sequence identity | Template PDBID | LogAUC for all the ligands | LogAUC for high affinity ligands |
| AA2AR | 4EIY | 27.37 | 30.08 | 20-30% | 3UON | 19.12 | 18.15 |
|  | 3REY | 23.00 | 23.48 |  | 5DSG | 20.11 | 20.16 |
|  | 2YDV | 25.08 | 25.67 |  | 2X72 | 25.09 | 24.31 |
|  | 4UHR | 28.23 | 30.65 |  | 4IB4 | 25.70 | 25.06 |
|  |  |  |  | 30-50% | 2VT4 | 19.34 | 18.94 |
|  |  |  |  |  | 3PBL | 17.14 | 18.26 |
|  |  |  |  |  | 2Y00 | 22.62 | 22.22 |
|  |  |  |  |  | 4LDE | 27.27 | 30.11 |
| ADRB2 | 2RH1 | 24.76 | 28.55 | 20-30% | 5CXV | 12.20 | 19.32 |
|  | 3NYA | 22.88 | 34.61 |  | 2Z73 | 9.27 | 12.80 |
|  | 4LDL | 23.27 | 24.63 |  | 4IB4 | 18.85 | 10.36 |
|  | 4LDE | 22.18 | 21.85 |  | 2X72 | 13.31 | 17.74 |
|  |  |  |  | 30-50% | 3PBL | 29.11 | 24.17 |
|  |  |  |  |  | 4U15 | 15.73 | 18.79 |
|  |  |  |  |  | 4IAR | 17.76 | 17.37 |
|  |  |  |  |  | 2YDO | 22.75 | 18.72 |
|  |  |  |  | 50-80% | 2VT4 | 22.06 | 21.63 |
|  |  |  |  |  | 4BVN | 19.27 | 25.58 |
|  |  |  |  |  | 5A8E | 23.88 | 26.49 |
|  |  |  |  |  | 2Y00 | 17.94 | 11.58 |
| LogAUC values of high affinity ligands are marked in bold to indicate significant improvement ($\Delta\text{logAUC} > 3$), and in italic to indicate comparable performance ($\vert \Delta\text{logAUC}\vert < 3$) | | | | | | | |

### Agonist-antagonist selectivity

To understand if an agonist-bound or antagonist-bounded X-ray structure or a homology model based on such a template confers any selectivity for either agonist or antagonist, a t-test was performed to check for any significant difference between the docking energies for agonists and antagonists (Table 8). This is because the difference in the average energy may seem significant, however it may still be a consequence of a few outliers and thus the data would be better represented through a statistical analysis.

**Table 8.** Average docking energy from agonist-bound and antagonist-bound X-ray structures and homology models

|  |  |  |  |  |  |  |
| --- | --- | --- | --- | --- | --- | --- |
| Receptor | PDBID | Cognate Ligand type | Agonist average docking energy | Antagonist average docking energy | t test P-value | Favour |
| AA2AR | 4EIY | Antagonist | -37.85 | -35.47 | 0.53 | - |
|  | 3REY | Antagonist | -35.13 | -28.07 | 0.00 | **Agonist** |
|  | 2YDV | Agonist | -40.77 | -27.82 | 0.22 | - |
|  | 4UHR | Agonist | -38.85 | -34.33 | 0.31 | - |
| Template sequence identity 20-30% | 3UON | Antagonist | -28.25 | -27.38 | 0.72 | - |
|  | 5DSG | Antagonist | -6.25 | -12.47 | 0.71 | - |
|  | 2X72 | Agonist | -15.41 | -14.98 | 0.98 | - |
|  | 4IB4 | Agonist | -31.84 | -27.63 | 0.11 | - |
| Template sequence identity 30-50% | 2VT4 | Antagonist | -32.45 | -28.11 | 0.11 | - |
|  | 3PBL | Antagonist | -37.85 | -32.78 | 0.21 | - |
|  | 2Y00 | Agonist | -27.77 | -25.44 | 0.33 | - |
|  | 4LDE | Agonist | -35.97 | -34.85 | 0.75 | - |
|  | 3D4S | Inverse agonist | -37.46 | -41.72 | 0.17 | - |
|  | 3SN6 | Agonist | -42.40 | -48.56 | 0.02 | **Antagonist** |
| ADRB2 | 2RH1 | Inverse agonist | -42.49 | -53.76 | 0.01 | **Antagonist** |
|  | 3NYA | Antagonist | -41.10 | -52.21 | 0.02 | **Antagonist** |
|  | 4LDL | Agonist | -47.30 | -49.27 | 0.63 | - |
|  | 4LDE | Agonist | -49.27 | -47.46 | 0.66 | - |
| Template sequence identity 20-30% | 5CXV | Antagonist | -37.58 | -40.40 | 0.62 | - |
|  | 2Z73 | Inverse agonist | -18.92 | -13.21 | 0.11 | - |
|  | 4IB4 | Agonist | -33.31 | -41.33 | 0.00 | **Antagonist** |
|  | 2X72 | Agonist | -51.26 | -57.79 | 0.12 | - |
| Template sequence identity 30-50% | 3PBL | Antagonist | -45.39 | -49.58 | 0.14 | - |
|  | 4U15 | Antagonist | -44.74 | -50.81 | 0.19 | - |
|  | 4IAR | Agonist | -35.08 | -49.16 | 0.00 | **Antagonist** |
|  | 2YDO | Agonist | -43.30 | -51.56 | 0.00 | **Antagonist** |
| Template sequence identity 50-80% | 2VT4 | Antagonist | -46.17 | -53.45 | 0.04 | **Antagonist** |
|  | 4BVN | Antagonist | -38.00 | -48.14 | 0.03 | **Antagonist** |
|  | 5A8E | Inverse agonist | -42.75 | -49.45 | 0.08 | - |
|  | 2Y00 | Partial agonist | -38.86 | -34.50 | 0.23 | - |

For AA2AR, there is almost no selectivity for either agonist or antagonist. This is expected as both agonists and antagonists have reasonable affinity to the protein and the docking energy function only considers the protein-ligand interaction energies. For ADRB2, all antagonist-bounded X-ray structures have significant tendency to select for antagonists over agonists while agonist-bounded X-ray structures do not show any selectivity. This is consistent with previous observations ^43, 44^. 5 out of the 12 homology models for ADRB2 shows selectivity for antagonists, among which almost equal number of them were built based on agonist-bound and antagonist-bound templates (3 and 2 respectively). One possible reason for this selectivity could be due to the fact that most X-ray structures of GPCRs were solved in their inactive states that prefer antagonists over agonists, no matter agonists or antagonists were used in the crystallization process. Another possible reason for this selectivity could be the chemical similarity between cognate ligands and the database agonists/antagonists. For ADRB2 X-ray structures, both cognate antagonists are similar to the database antagonists (average Tanimoto similarity > 0.4) and dissimilar to the database agonists. On the contrary, both cognate agonists show no significant similarity with either database agonists or antagonists. This likely contributed to the significant preference for antagonists by ADRB2 antagonist-bounded structures. It might also be possible that the observed selectivity by ADRB2 is a result of antagonists having higher affinity than agonists. Currently, there are insufficient information on the affinities of the agonists and antagonists to determine if that is the case For AA2AR, only 1 agonist have affinity value (Ki/Kd) thus making affinity comparison not feasible. For ADRB2, out of the 4 antagonists with affinity value (Ki/Kd), all of them are below 10nM while out of the 7 agonists with affinity value, only 2 of them are below 10nM. However, the difference are not statistically significant (p-value > 0.05) to make any conclusion.

## Conclusions

### Overview

While the majority of the homology models were outperformed by the X-ray structures in virtual screening, noticeably 15 out of the 38 homology models performed better than or comparable to the corresponding X-ray structures, and 30 out of the 38 models are value-add to virtual screening, indicating the usefulness homology models for virtual screening.

There is no clear correlation between the ligand enrichment (logAUC) from homology models and their sequence identity to templates. However, it was noted that homology models with 50-80% sequence identity have similar chance to value-add to virtual screening as the X-ray structures, exhibiting the influence of sequence identity level on the virtual screening performance of homology models.

### Consensus over multiple homology models

Consensus enrichment scores over multiple homology models based on different templates, calculated by selecting the best docking score for each compound, can perform consistently close to the top performing homology model. Thus, when the quality of homology models in ligand recognition is unclear, consensus models are recommended over multiple models.

### Alternative sidechain conformations

For X-ray structures, sidechain conformation optimization has no significant improvement to virtual screening, which needs to be further evaluated before a conclusion can be made. For homology models, there is clear indication that the ligand recognition ability of one model can be significantly improved by sidechain reconstruction in the presence of a known ligand.

### High affinity ligands

Homology models had no significant tendency to select high affinity ligands over lower affinity ligands in virtual screening, indicating that the structure-based docking approach may not be able to accurately differentiate ligands of different affinities.

### Agonist-antagonist selectivity

There is some indication that for certain receptor such as ADRB2, that their antagonist-bound or inverse agonist-bound X-ray structures have higher selectivity for antagonists over agonists; while for homology models, the preference for antagonists can be observed from models based on both agonist-bound and antagonist-bound templates.

In conclusion, our studies suggest the following strategies for exploiting homology models of GPCRs in virtual screening: 1) when multiple templates are available, homology models of GPCRs are best used *via* consensus ligand selection including multiple models based on different templates; 2) when there is only one template available, the resulted model can be further improved for ligand recognition by local conformational sampling in the presence of known ligands.

## Acknowledgement

The work was supported by Biomedical Research Council of A*STAR (to Hao Fan and Weina Du), and by the Singapore Academic Research Fund R-148-000-230-112 and R-148-000-239-112 (to Yu Zong Chen)

## Abbreviation list

5HT1B: 5-Hydroxytryptamine Receptor 1B
5HT2B: 5-Hydroxytryptamine Receptor 2B
AA2AR: Adenosine A2a Receptor
ACM2: Muscarinic Acetylcholine Receptor M2
ACM3: Muscarinic Acetylcholine Receptor M3
ADRB1: Beta-1 Adrenergic Receptor
ADRB2: Beta-2 Adrenergic Receptor
CCR5: C-C Chemokine Receptor Type 5
CRFR1: Corticotropin-releasing factor receptor 1
CXCR4: C-X-C Chemokine Receptor Type 4
DRD3: Dopamine Receptor D3
GCCR: Glucagon Receptor
GPR40: G-protein-coupled Receptor 40/ Free Fatty Acid Receptor 1
HRH1: Histamine Receptor H1
MGLUR1: Metabotropic Glutamate Receptor 1
MGLUR5: Metabotropic Glutamate Receptor 1
OPRD: Opioid Receptor Delta 1
OPRK: Opioid Receptor Kappa 1
OPRM: Opioid Receptor Mu 1
OPRX: N/OFQ Opioid Receptor
OX2R: Orexin Receptor Type 2
P2Y12: Purinergic Receptor P2Y12
PAR1: Protease-activated Receptor 1
S1PR1: Sphingosine-1-phosphate Receptor 1
AT1R: Angiotensin Type 1 Receptor
NTSR1: Neurotensin Receptor 1
LPAR1: Lysophosphatidic Acid Receptor 1
RHO: Rhodopsin

