## Supplementary Materials for "A benchmarking study on virtual ligand screening against homology models of human GPCRs"

### <u>Supporting information</u>

**Content of supporting information**

Table S1. Table of X-ray structures and templates for AA2AR and ADRB2.

Table S2. List of high affinity ligands for AA2AR and ADRB2

Table S3. List of agonists and antagonists of AA2AR and ADRB2.

Table S4. Ligand enrichment of X-ray structures with predicted sidechains

Figure S1. Histogram of docking energy distribution of high affinity ligand (grey) and low affinity ligands (black).

**Table S1. X-ray stuctures of AA2AR and ADRB2 and their templates.**

| GPCR | Ligand Type | X-ray structure | Template 20-30% | | | Template 30-50% | | | Template 50-100% | | |
|---|---|---|---|---|---|---|---|---|---|---|---|
| | | PDBID | Sequence identity% | PDBID | Template GPCR | Sequence identity% | PDBID | Template GPCR | Sequence identity% | PDBID | Template GPCR |
| AA2AR | Antagonist/ Inverse Agonist | 4EIY | 25 | 3UON | ACM2 | 36 | 2VT4 | ADRB1 | | | |
| | | 3REY | 27 | 5DSG | ACM4 | 35 | 3PBL | DRD3 | | | |
| | Agonist/ Partial Agonist | 2YDV | 23 | 2X72 | RHO | 35 | 2Y00 | ADRB1 | | | |
| | | 4UHR | 27 | 4IB4 | 5HT2B | 31 | 4LDE | ADRB2 | | | |
| ADRB2 | Antagonist/ Inverse Agonist | 2RH1 | 28 | 5CXV | ACM1 | 36 | 3PBL | DRD3 | 58 | 2VT4 | ADRB1 |
| | | 3NYA | 24 | 2Z73 | RHO | 31 | 4U15 | ACM3 | 58 | 4BVN | ADRB1 |
| | Agonist/ Partial Agonist | 4LDL | 27 | 4IB4 | 5HT2B | 33 | 4IAR | 5HT1B | 57 | 5A8E | ADRB1 |
| | | 4LDE | 22 | 2X72 | RHO | 30 | 2YDO | AA2AR | 58 | 2Y00 | ADRB1 |

| Table S2. ChEMBL ID of high affinity ligands for AA2AR and ADRB2 | | |
|---|---|---|
| AA2AR | | ADRB2 |
| CHEMBL399577 | CHEMBL475445 | CHEMBL1221736 |
| CHEMBL259442 | CHEMBL16687 | CHEMBL2088201 |
| CHEMBL193761 | CHEMBL487553 | CHEMBL2426777 |
| CHEMBL3100164 | CHEMBL1800358 | CHEMBL266195 |
| CHEMBL429428 | CHEMBL113142 | CHEMBL513389 |
| CHEMBL360241 | CHEMBL255911 | CHEMBL183921 |
| CHEMBL1762506 | CHEMBL409682 | CHEMBL1083668 |
| CHEMBL392136 | CHEMBL270880 | CHEMBL1085010 |
| CHEMBL471853 | CHEMBL482161 | CHEMBL251392 |
| CHEMBL112631 | CHEMBL323945 | CHEMBL513104 |
| CHEMBL456287 | CHEMBL550636 | CHEMBL1221587 |
| CHEMBL1935760 | CHEMBL2377233 | CHEMBL393648 |
| CHEMBL260718 | CHEMBL2419149 | CHEMBL1221802 |
| CHEMBL2030679 | CHEMBL373026 | CHEMBL1233771 |
| CHEMBL1096998 | CHEMBL2030690 | CHEMBL36060 |
| CHEMBL3121726 | CHEMBL2165804 | CHEMBL1221590 |
| CHEMBL3121721 | CHEMBL2377110 | CHEMBL1221543 |
| CHEMBL1086847 | CHEMBL3121731 | CHEMBL1221803 |
| CHEMBL402071 | CHEMBL260719 | CHEMBL1221681 |
| CHEMBL1672619 | CHEMBL413079 | CHEMBL1221634 |
| CHEMBL486552 | CHEMBL248298 | CHEMBL1221804 |
| CHEMBL2377103 | CHEMBL271202 | CHEMBL1221541 |
| CHEMBL372475 | CHEMBL168334 | CHEMBL1094785 |
| CHEMBL166429 | CHEMBL1095999 | CHEMBL122990 |
| CHEMBL443665 | CHEMBL485972 | CHEMBL723 |
| CHEMBL594905 | CHEMBL259323 | CHEMBL499 |
| CHEMBL256123 | CHEMBL20888 | CHEMBL1221861 |
| CHEMBL27508 | CHEMBL1762511 | |
| CHEMBL519566 | CHEMBL1270115 | |
| CHEMBL2030687 | CHEMBL1093271 | |
| CHEMBL611335 | CHEMBL177782 | |
| CHEMBL511405 | CHEMBL144521 | |
| CHEMBL486745 | CHEMBL519549 | |
| CHEMBL145623 | CHEMBL1650384 | |
| CHEMBL359463 | CHEMBL272924 | |

| **Table S3. Agonists and antagonists of AA2AR and ADRB2.** | | | |
|---|---|---|---|
| AA2AR | | ADRB2 | |
| Agonist | Antagonist | Agonist | Antagonist |
| CHEMBL317052 | CHEMBL628 | CHEMBL500 | CHEMBL499 |
| CHEMBL1950554 | CHEMBL240624 | CHEMBL714 | CHEMBL649 |
| CHEMBL1950649 | CHEMBL2105747 | CHEMBL1263 | CHEMBL27 |
| CHEMBL1950553 | CHEMBL447664 | CHEMBL1198857 | CHEMBL471 |
| CHEMBL1950651 | CHEMBL197669 | CHEMBL1760 | CHEMBL1290 |
| CHEMBL3545163 | CHEMBL113 | CHEMBL434 | CHEMBL723 |
| CHEMBL477 | CHEMBL190 | CHEMBL679 | CHEMBL839 |
| CHEMBL499 | CHEMBL113142 | CHEMBL1546 | CHEMBL429 |
| CHEMBL281213 | CHEMBL2024115 | CHEMBL1201234 | CHEMBL266195 |
| | | CHEMBL1437 | CHEMBL1233771 |
| | | CHEMBL2103827 | CHEMBL324665 |

| **Table S4. Ligand enrichment of X-ray structures with predicted sidechains** | | | | | | | |
|---|---|---|---|---|---|---|---|
| Receptor | PDBID | X-ray structure | | SCWRL | | PLOP | |
| | | LogAUC | EF1 | LogAUC | EF1 | LogAUC | EF1 |
| AA2AR | 4EIY | 27.37 | 4.93 | 29.92 | 4.44 | 26.81 | 3.45 |
| ADRB2 | 2RH1 | 24.76 | 6.98 | 23.75 | 5.99 | 27.68 | 5.49 |

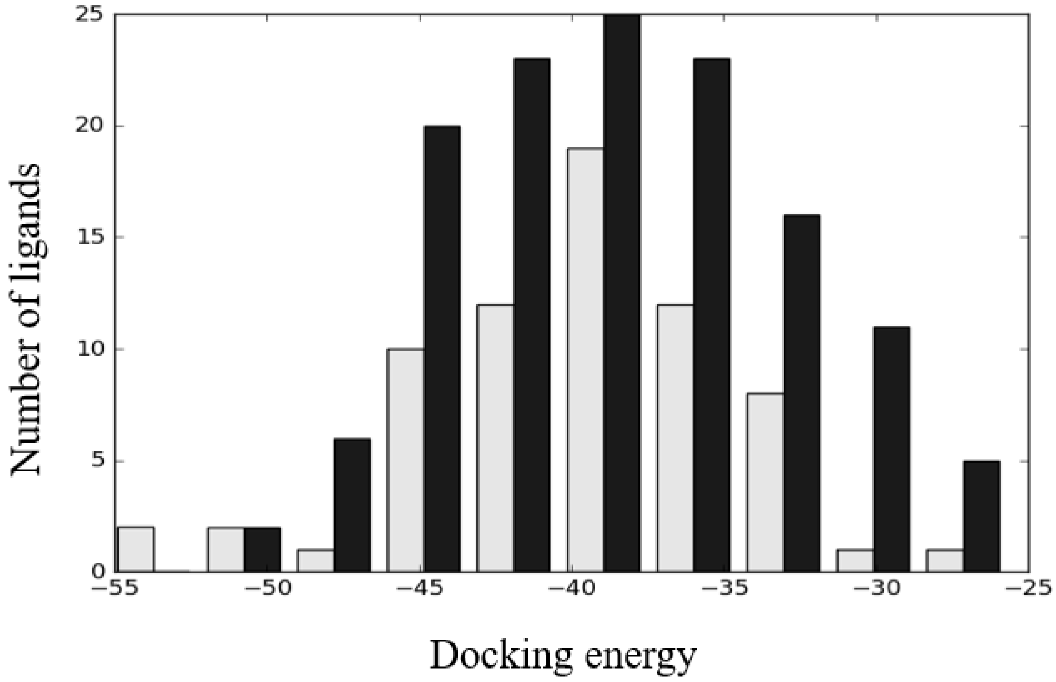


**Figure S1.** Histogram of docking energy distribution of high affinity ligand (grey) and low affinity ligands (black).
